# Early *in planta* detection of *Xanthomonas axonopodis* pv. *punicae* in pomegranate using enhanced loop-mediated isothermal amplification assay

**DOI:** 10.1101/328674

**Authors:** M.K. PrasannaKumar, P. Buela Parivallal, C. Manjunath, H.B. Mahesh, Karthik S Narayan, Gopal Venkatesh Babu, K. Priyanka, M.E. Puneeth, K.T. Rangaswamy

## Abstract

Bacterial blight in pomegranate caused by *Xanthomonas axonopodis* pv. *punicae* (*Xap*) is an increasing threat for pomegranate cultivation in India. To prevent the economic losses, it is pivotal to detect the infection in latent stages rather than in later stages. We have developed an enhanced method termed as loop-mediated isothermal amplification (LAMP) technique to evaluate for the latent detection of Xap in pomegranate using six set of specific primers. Three DNA intercalating dyes were used, such as Ethidium bromide, hydroxynaphthol blue (HNB) and SYBR Green resulted in visualising the positivity for LAMP assay. The reaction time and temperature were to be 65°C from 30 min onwards, for the dyes and its sensitivity was observed up to 10^−7^ ng in the LAMP assay. For field applicability, LAMP assay detected Xap on 7^th^ day post infection while the PCR amplified Xap after 11^th^ day post infection. Finally, the specificity of LAMP assay was validated to be positive with ten Xap isolates for its accuracy and 29 non-Xap bacterial isolates showed negative results. Moreover, this method could be used as a better alternative to PCR based methods, for early detection of the pathogens.

## Introduction

Pomegranate (*Punica granatum* L.) is an important fruit crop widely cultivated from tropical to temperate conditions [1]. India ranks first in the world for maximum area under pomegranate cultivation (0.125 million hectares) and Maharashtra, Karnataka and Andhra Pradesh are the major pomegranate growing states [2]. The bacterial blight caused by *Xanthomonas axonopodis* pv. *punicae* (Xap) in pomegranate was first reported in India in 1952 and presently it has a serious outbreak in all major pomegranate-growing states resulting in a major economic loss. Prominent small water-soaked and dark irregular spots appearing on the leaf are the typical symptoms induced by this pathogen. The pathogen infect the fruit, stem, branches causing cracking symptoms and ultimately leads to yield loss up to 80 percent [3].

Globally, it is estimated that a minimum 30-40% of the crop cultivated gets destroyed annually due to pests and diseases [4]. In India, bacterial blight alone reduces the yield between 60 to 80 percent above global yield loss in pomegranate. To avoid yield loss, it is very important and necessary to develop an efficiently sensitive pathogen detection method within incubation time of the pathogen (i.e. time between the host infection and expression of the disease symptoms) [5,6].

The most common methods employed for detecting the plant pathogens include immunological techniques like ELISA (enzyme-linked immunosorbent assay) [7] and molecular methods such as PCR (polymerase chain reaction) [8] and Southern blotting [9]. The methods for early detection of the plant pathogens in the field has to be designed in such a way that it has to be competently very sensitive, specific, cost-effective and detectable with a minimal equipment facilities [10]. For instance, the PCR based technique has its own drawbacks which require high quality and quantity of DNA and amplification is complex that requires a thermal cycler for changing the reaction temperature and requires relatively long time for the amplification process to complete. Further, it also requires agarose gel electrophoresis and gel documentation to confirm the amplification [11]. To overcome these limitations, researchers have shown interest in developing an isothermal amplification technique termed as Loop-mediated isothermal amplification (LAMP) [12]. LAMP is an auto-cycling strand displacement DNA synthesis technique, which is carried out at a constant temperature and does not require a thermal-cycler. The amplification is completed within 30-60 minutes at a constant temperature of 60-65° C, which can be carried out in a simple water bath or heating block [13]. Moreover, LAMP is a cost effective and it does not require a thermal cycler and electrophoretic unit, as the result can be inferred as positive or negative with the colour change or turbidity observed by naked eye without spending much time [14].

The favorable target for bacterial pathogen diagnostic includes mainly, 16S ribosomal RNA (rRNA) gene which is been widely used to identify the bacterial species among different bacteria [15–17]. This diversity is unique for species and their sequences are presented inside their nine (V1 - V9) hypervariable regions which constitutes useful targets [18]. These regions are highly conserved stretches in bacteria, which facilitates the identification through PCR amplification using universal primers [19].

In principle, the LAMP assay recognizes six specifically designed primers that recognize eight distinct sequences from the target bacterial DNA [20]. It consists of two outer primers (F3 and B3), two inner primers (FIP and BIP), a loop-forward (LF) and a loop-backward (LB). These six primers ensure a high specificity for target DNA amplification. Moreover, these primers enable generation of a stem-loop DNA for subsequent complex LAMP amplification, which includes self-priming reaction. A mixture of DNA, differing in the number of loops and length of the stem-loop is produced as a final product [21]. By considering the advantages of LAMP assay, which is widely used to detect broad range of microorganisms[22,23]. The present study was aimed to detect the bacterial blight of pomegranate in early stages with minimum pathogen DNA template by optimizing several DNA intercalating dyes with lower temperature in a shorter time with less technical skills.

## MATERIALS AND METHODS

### Source of bacterial blight material used for the study

Ten *Xanthomonas axonopodis* pv. *punicae* (Xap) isolates collected from four different regions of Karnataka, India were used in this study (Fig 1). Additionally, we have also isolated 29 non-Xanthomonas bacteria from pomegranate Table 1. The pure culture Xap isolates were maintained in a Nutrient Broth for 72 hours and the DNA was extracted using crude method. The quality and quantity of the DNA were determined using Nanodrop (Model - DS-11 FX +, DeNovix, USA).

**Figure 1.**
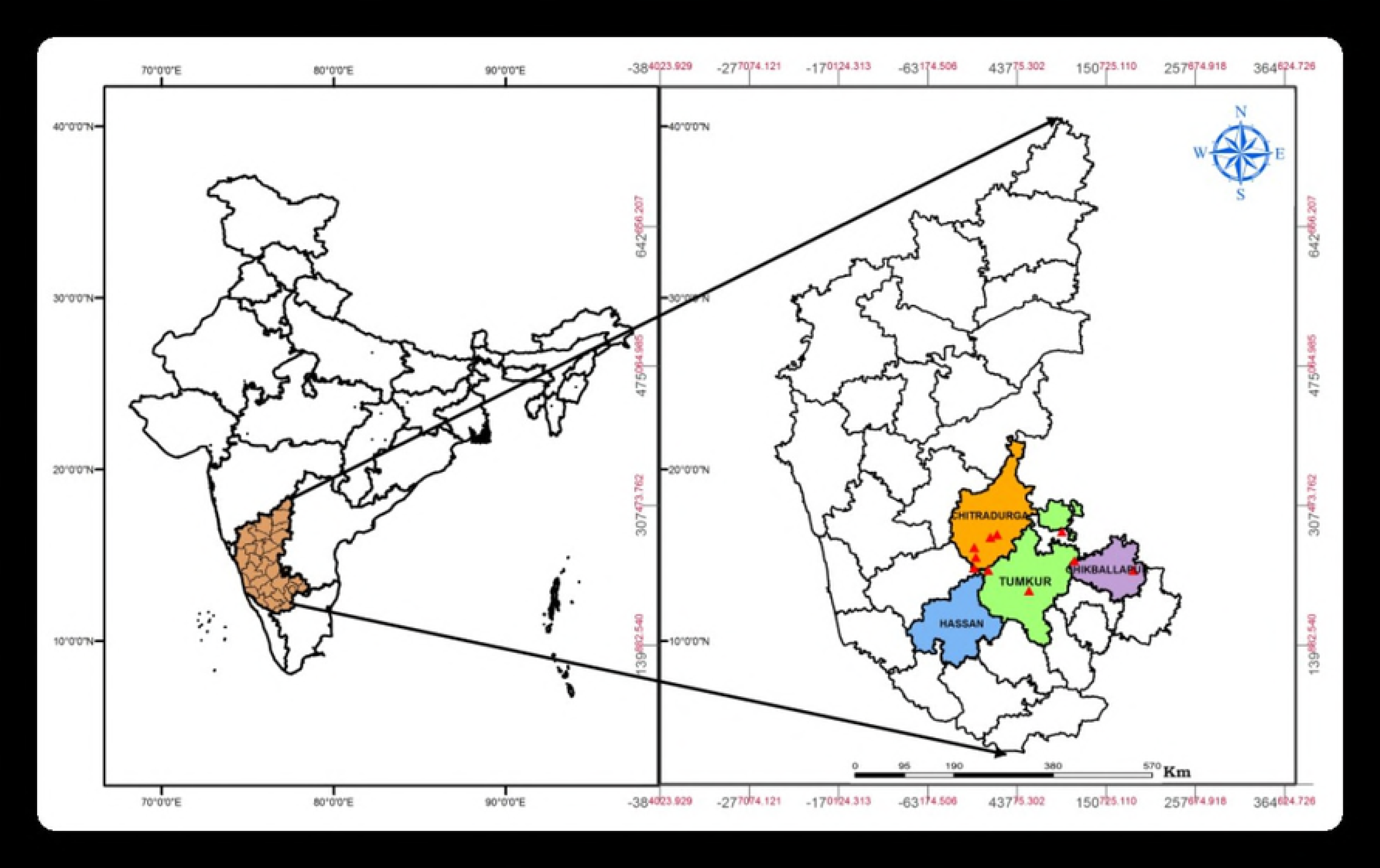
GIS mapping of collection sites of Xap and other bacteria isolates from Karnataka, India. The ten sampling points were shown within four districts of Karnataka namely Hassan, Tumkur, Chitradurga and Chikkaballapur respectively.

**Table 1.**
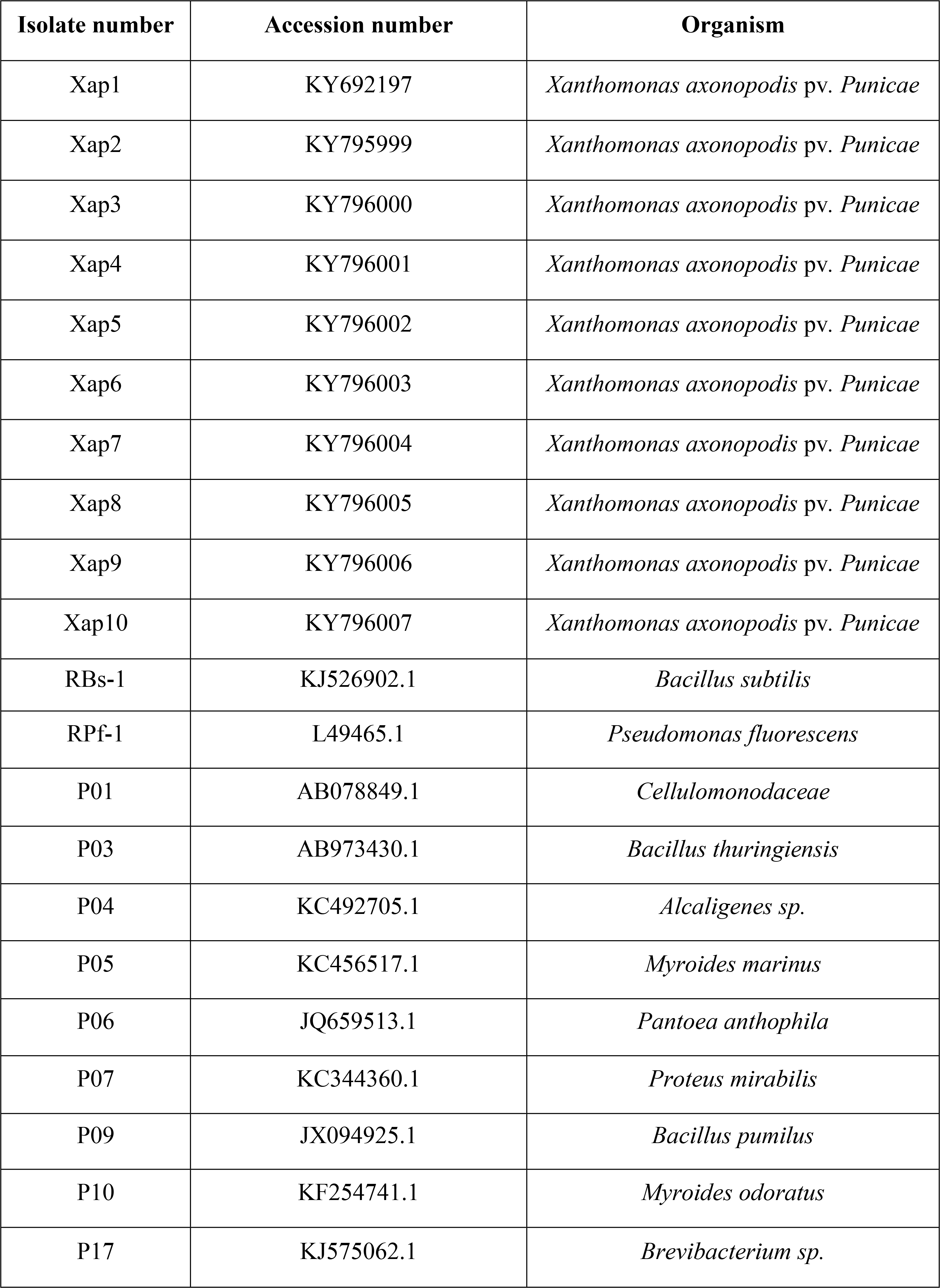
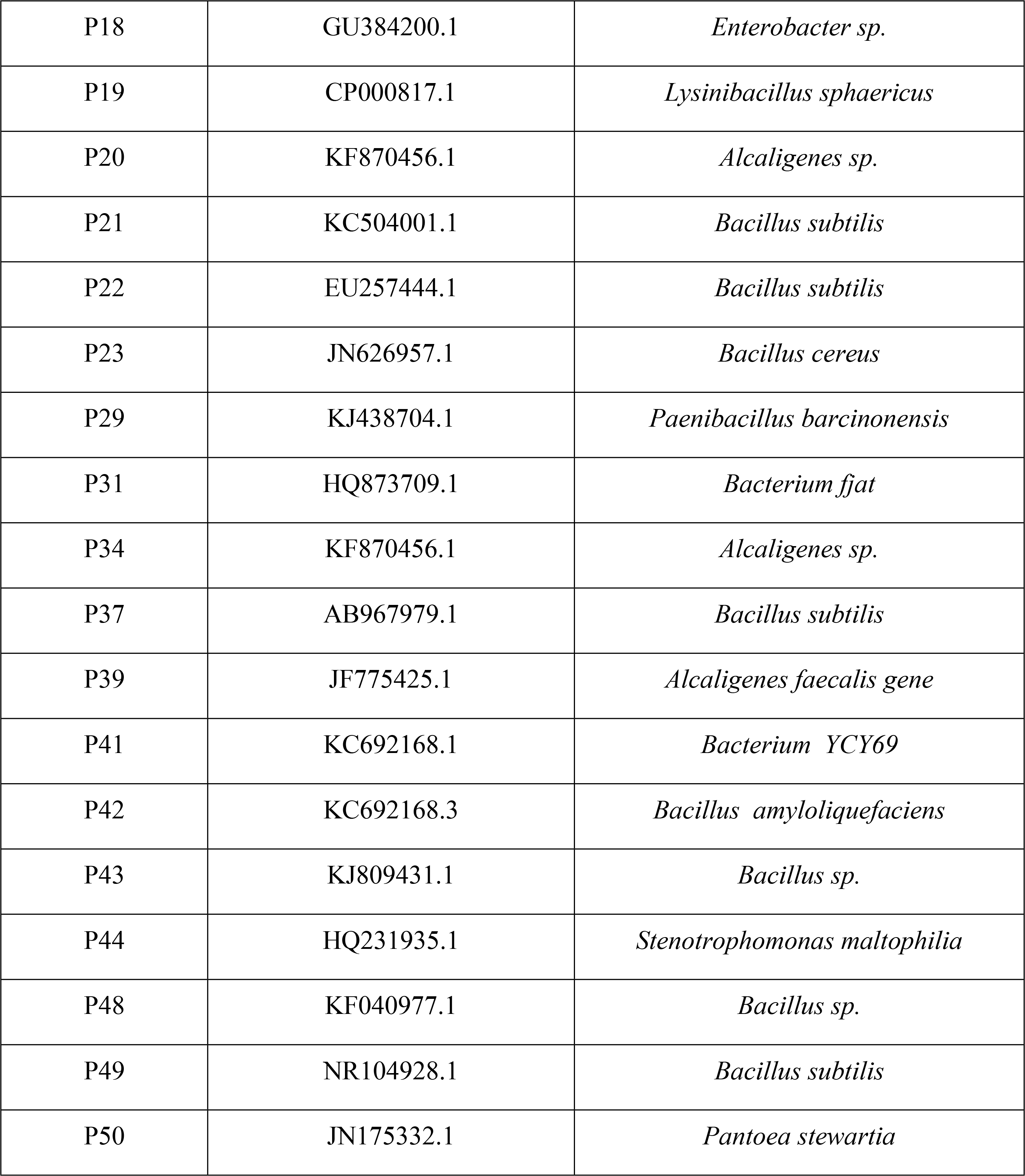
Details of Xap and non-Xap bacterial isolates used for specificity assay. All the collected isolates checked for amplification of Xap gene in both PCR and LAMP assays.

### Crude method

To test the cost effectivity and latent detection of bacterial blight pathogen, the DNA was isolated from the leaf sample collected from glass house, using a modified crude method [24]without the use of centrifuge and other instruments. The crude extract was used as a template for LAMP assay as well as PCR reaction setup. The leaf sample collected from glass house was surface sterilized with 0.1% mercuric chloride for about 30 to 90 seconds, later the leaf was washed thrice with sterile water to remove any residues of mercuric chloride. The infected plants tissue were immersed in the tube containing 500 μL of STE buffer (100 mM NaCl, 100 mM Tris and 50 mM EDTA) for 10 minutes and similarly for other tubes 50 μLof pure cultures (Xap and other bacterial non Xap spp.) was added. The tubes were vigorously mixed to form the uniform solution of bacteria, 500 μL of phenol: chloroform: isoamyl alcohol mixture (25:24:1) was added and mixed vigorously and allowed to stand till the aqueous and organic phase separated. Then the supernatant was collected (containing genomic DNA) into new sterile tube for further analysis.

### Primer design

All the DNA from ten different Xap isolates were used to sequence16s rRNA region using specific primer reported by Mondal *et al.*, [25] (Forward 5’-AGAGTTTGATCCTGGCTAG-3’ and Reverse 5’-AGGAGGTGATCCAGCCGCA-3’The LAMP assay encodes for six specifically designed primers that recognize eight distinct sequences from the sequenced amplified product of 16S rRNA from Xap 1. A set of six primers for LAMP assay, comprising two each of outer, inner and loop primers were designed by Primer Explorer V4 http://primerexplorer.jp/e, Japan; Table 2). The position of primers and their complementary to its target sequences are shown diagrammatically in Fig 2. PCR based amplification was carried out using Xap specific primer set, which were designed from the 16s rRNA sequence (Forward-5’-CTGGAAAGTTCCGTGGAT-3’; Reverse-5’-TTGCAGTGGATACTGGGT-3’).

**Table 2.**
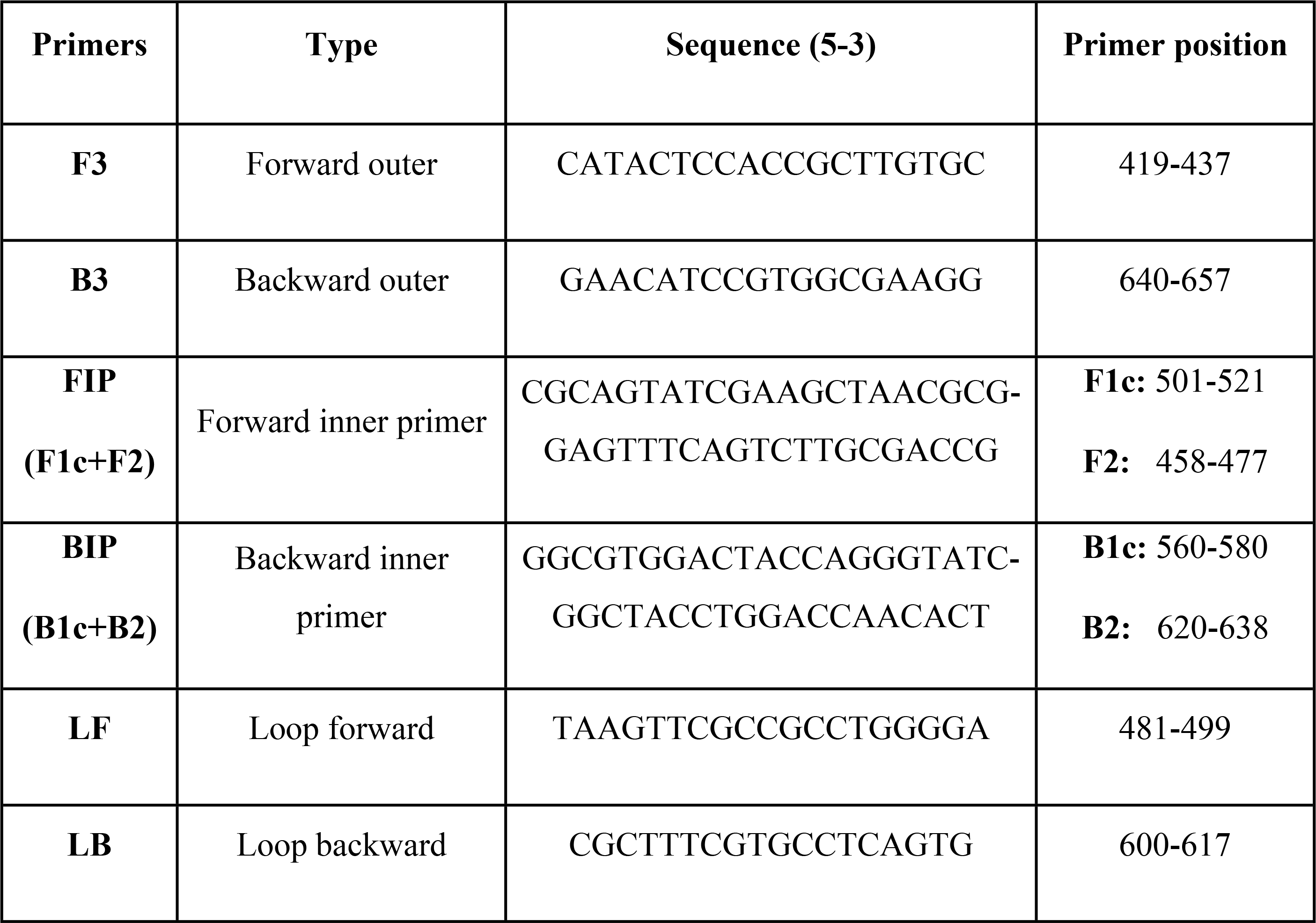
Oligonucleotides used for the detection of *Xanthomonas axonopodis* pv. *punicae* for LAMP assay. The Forward inner primer consists of two regions namely F1c and F2 likewise; the backward inner primer consists of B1c and B2 regions.

**Figure 2.**
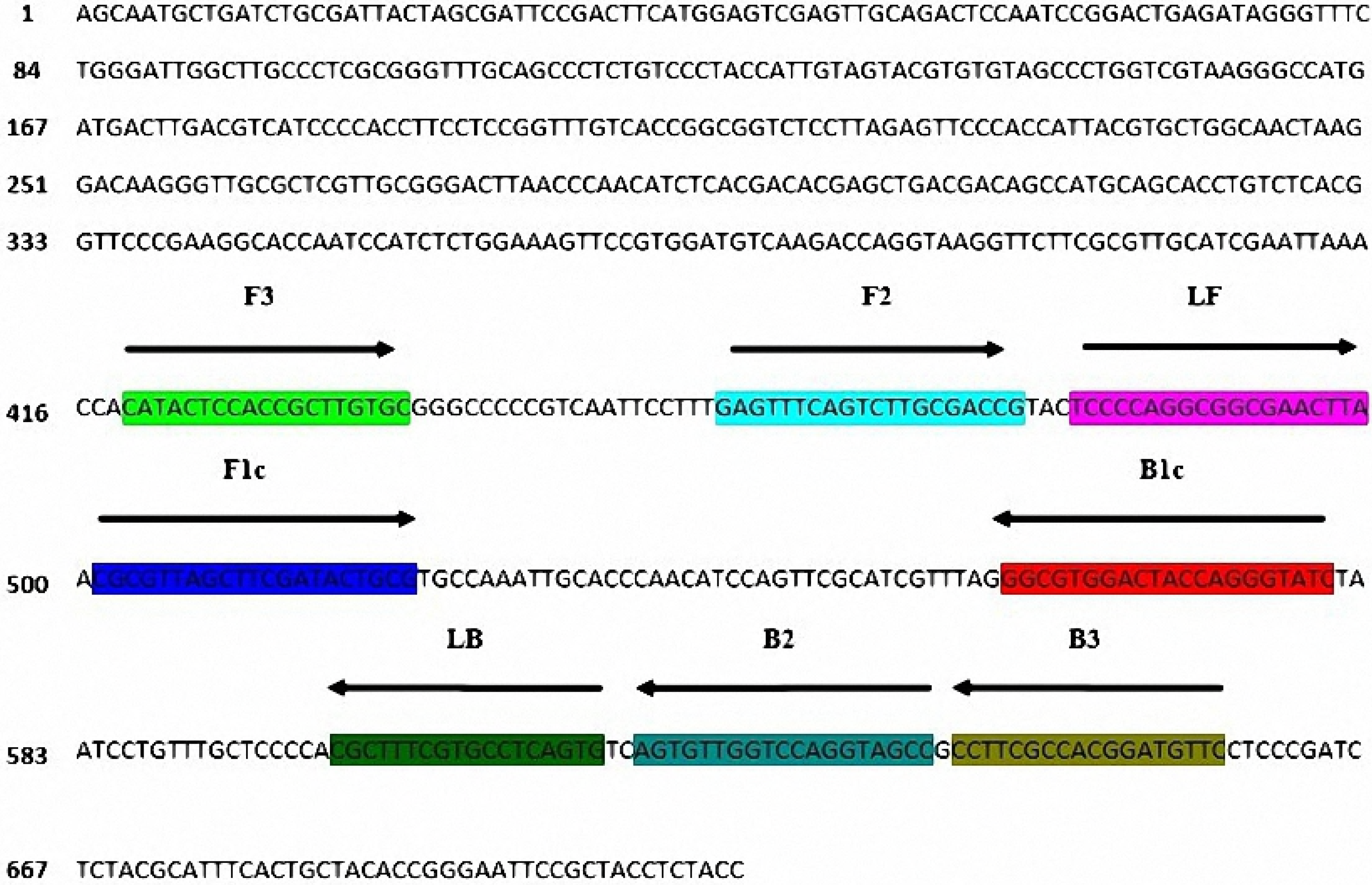
Location and sequence of LAMP primers specifically designed for Xap. The location of six primers [F3, B3, FIP (F1c+F2), BIP (B1c+B2), LF and LB] and arrows indicating the direction of extension. Detailed explanation of the primers are given in table 2.

### LAMP and PCR reaction setup

LAMP assay was carried out in a 25 μL reaction mixture containing of 1X Thermo pol buffer pH 8.8 (10 mM (NH_4_)_2_SO_4_, 20 mM Tris-HCl, 10 mM KCl, 2 mM MgSO_4_, 0.1% Triton-X), 0.2 M betaine, 0.6 mM of dNTPs, 10 pmol of F3 (forward outer) and B3 (backward outer) primers, 20 pmol of FIP (forward inner primer), BIP (backward inner primer), LF (loop forward) and LB (loop backward) primers, 8 U of *Bst* polymerase large fragment, template DNA with a concentration of 1.53 ng/ μL and the reaction volume was made up to 25 μL using sterile water. The reaction volume and the setup were mentioned in Table S1. The reaction mixture was incubated at 65° C for 1 hour and the process was terminated at 80° C for 10 mins using a hot water bath (KEMI, India). Further for more visual differentiation to confirm the LAMP results (positive or negative), a total of 5 DNA intercalating dyes were used and their details were presented in table 3. The emission (RFU) was measured using Basic Fluorometer for the dyes EtBr and SYBR Green which was measured using UV Excitation at ~375nm in a Basic fluorometer and for HNB dye the blue excitation was measured at ~470nm in a Nanodrop (Model - DS-11 FX +, DeNovix, USA). The amplified product was subjected to agarose gel electrophoresis on a 2% agarose gel and documented in a software gel documentation system (Model - Molecular Imager Gel Doc XR+, BioRad, USA and software - Version 5.2.1). The illustration of the LAMP assay was described in pictorial form Fig 3.

**Table 3.** Details of the DNA intercalating dyes. A total of five intercalating dyes used in the LAMP assay and its field applicability were described and indicated as positive (+) or negative (−).

|  | **Ethidium bromide** | **SYBR Green** | **Hydroxy naphthol blue (HNB)** |
| --- | --- | --- | --- |
| **Stock concentration** | 10 mg/mL | 1 X | 20 mM |
| **Volume per (µL)** | 1.0 | 0.5 | 1.0 |
| **Treatment** | Post-treatment | Post-treatment | Pre-treatment |
| **Detection system** | UV | UV | Visualised through naked eyes |
| **Expected Result** | + Bright Orange<br>- Light Orange | + Green<br>- Transparent | + Blue<br>- Purple |
| **Field applicability** | + | + | + |

**Figure 3.**
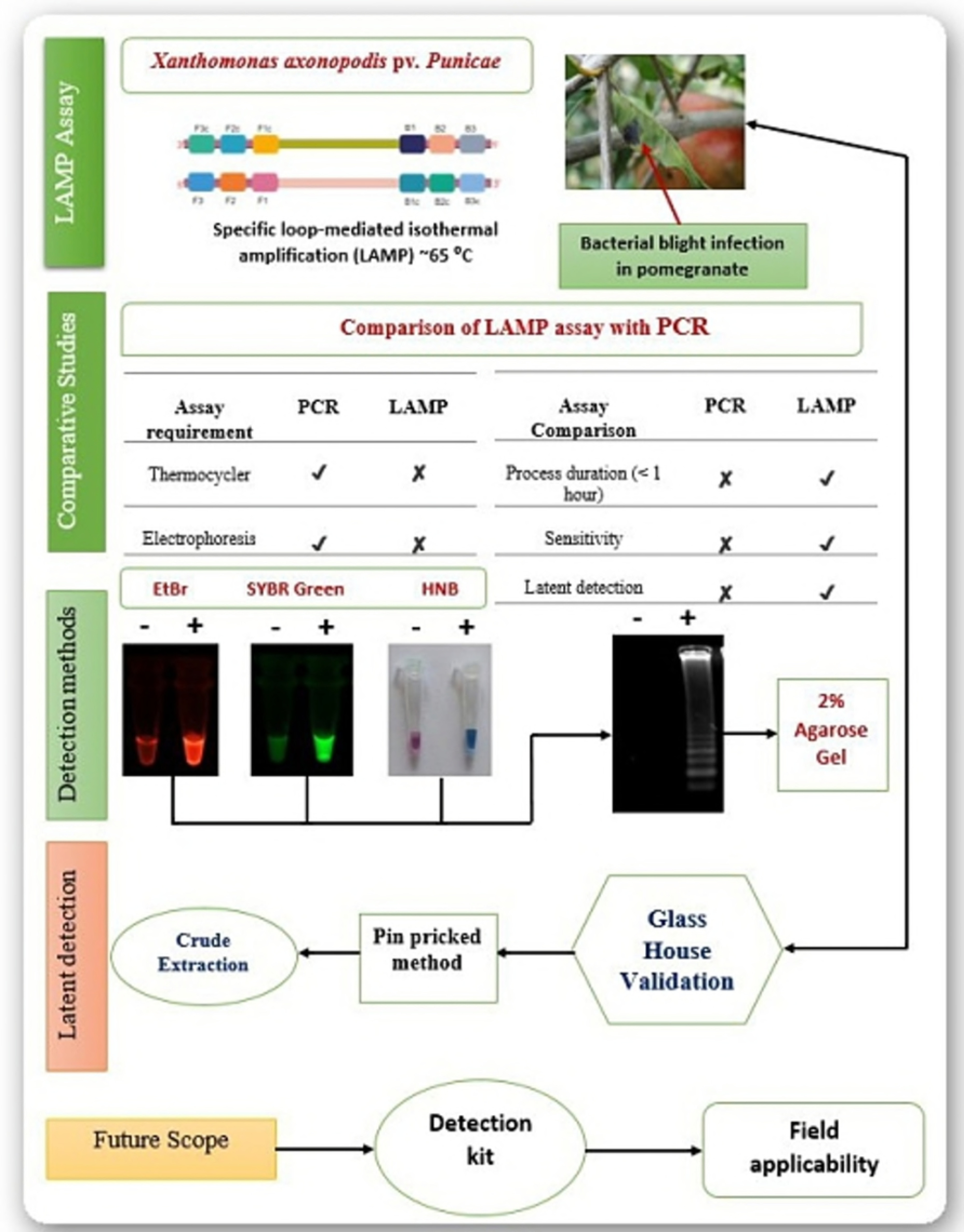
Overview of LAMP assay. The PCR reaction was carried out in a thermal cycler (Eppendorf- vapo.protect, Germany) with the following conditions; Initial denaturation at 94°C for 5 min, followed by 30 cycles of denaturation at 94°C for 30 seconds, annealing at 55°C for 60 seconds and extension at 72°C for 176 1 min. At the end of 35 cycles, the final extension was at 72°C for 3 min. The amplified products were visualised on 1% agarose gel electrophoresis.

### Optimization and Sensitivity of LAMP assay

The LAMP reactions were carried out to optimize a single temperature for greater efficiency. The reaction was performed at different temperatures ranging from 60 to 65°C. After which the time duration was optimized from 0 to 90 min. The initial concentration of the template DNA of the isolate was adjusted to 1.53 ng/μL for serial diluted samples in a series of 10 fold series (10^−1^ to 10^−10^ ng/μL) that was carried to determine the sensitivity of LAMP assay for different DNA intercalating dyes Table 3. The sensitivity of PCR and LAMP assay was compared using different concentrations of template DNA in the reaction mixtures.

### Validation for specificity

It is very important to validate the LAMP assay and its dyes with respect to its specificity (only Xap) and accuracy (among the Xap samples). To analyze the accuracy, we have used 10 isolates of Xap and 29 non Xap isolates of bacteria isolated from the same source, pomegranate. Isolates with each dye were compared using completely randomized block design to identify the significant difference by Pairwise Mean Comparison on the variants using Duncan’s Multiple Range Test (DMRT). Moreover, each isolate was checked with all the dyes for the ten isolates using one sample ‘t’ test. Statistical tool (R version 3.4.3) was used to find the significant differences in both DMRT and one sample ‘t’ test.

### Latent *in planta* detection of Xap

The detection studies were performed in a controlled humidity and temperature conditions in the glass house at University of Agricultural Sciences, G.K.V.K, Bangalore, Karnataka, India (Department of Plant Pathology) for validating latent detection between LAMP and PCR methods. Three months old pomegranate plant (Bhagwa va.) was infected by pin pricked method [26]. The Xap was suspended (1 × 10^8^ CFU/mL) in 0.01 M phosphate buffer (pH 7.2) and sterile needle was used to prick the leaves. The plants were covered with a polythene cover along with water sprinkled in order to maintain the humidity. The leaf samples were collected after the inoculation at alternative days.

## Results

### Optimization of LAMP assay using different dyes

A total of 10 Xap isolates (Xap1 to Xap10) were used to optimize the LAMP assay. Colour differentiation between positive and negative assays for the detection of Xap was the main criteria for selection of different dyes. Three DNA intercalating dyes (Ethidium bromide, Hydroxynaphthol blue and SYBR Green) were used in the LAMP assay. The primers were designed from the sequence of 16S rRNA of Xap1 isolate. The dyes EtBr, HNB, and SYBR green were found to be optimum for LAMP assay based on their feasibility in detection Table 3. To determine the optimum temperature and reaction time of LAMP assay, a short range of temperature (60°C - 65°C) was used to analyse the highest fluorescent intensity and time (15 min - 90 min) using high quality DNA of Xap1 isolate. LAMP assay was positive for all the three dyes (EtBr, HNB, SYBR Green) used with respect to various levels of temperature (Fig 4a). EtBr showed the highest fluorescent intensity with increase in temperature compared to other dyes (Fig 4b) which was also assessed on 2% agarose gel (Fig 4c) and the fluorescent intensity was also measured simultaneously. Amplification was not observed at 15 min incubation time and positive LAMP results were observed 30 min onwards (Fig 4d) and further it was reconfirmed on 2% agarose gel (Fig 4e). The highest fluorescent intensity was recorded at 75 min (Fig 4f) for each dye. Therefore, Xap can be best detected using EtBr under an optimized condition at 65° C from 30 minutes onwards under UV light. The fluorescent intensity was decreased to half of that of EtBr when the assay was directly visualised using HNB dye without UV-detector.

**Figure 4.**
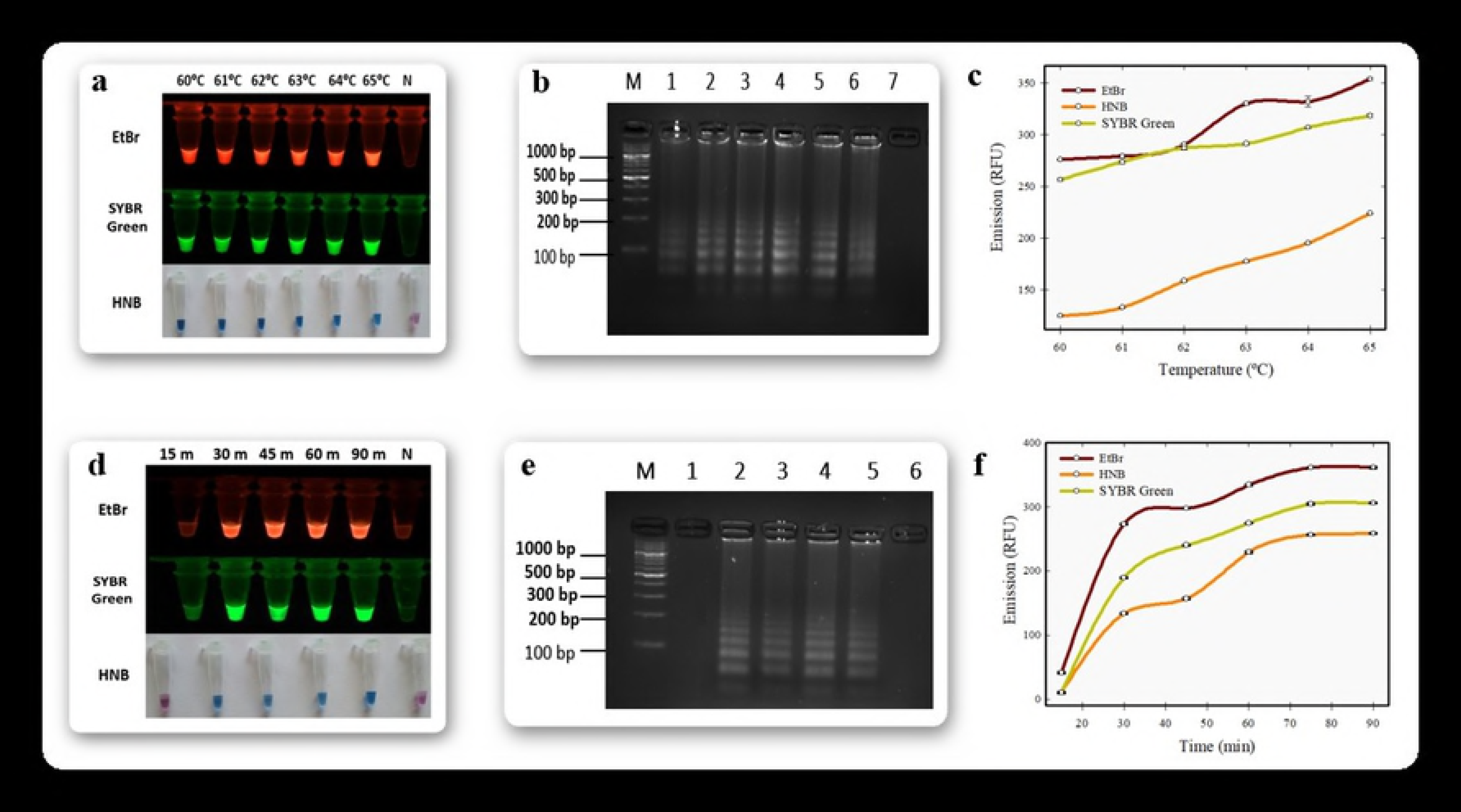
Optimization of temperature and time for the LAMP assay. (**a**) Temperature optimisation of LAMP with EtBr, SYBR Green and HNB dyes ranged between 60°C - 65°C. (**b**) Further, it was visualised in agarose gel Lane (1 to 6) indicated 60°C - 65°C and Lane N indicates negative control without a template. (**c**) Optimisation graph of LAMP with Temperature for each dyes. (**d**) Time optimisation of LAMP with EtBr, SYBR Green and HNB dyes ranged between 15 to 90 min. (**e**) Further, it was visualised in agarose gel, Lanes (1 to 5) indicated time (15, 30, 45, 60 and 90 min) and Lane N indicated negative control without any template. (**f**) Optimisation graph of LAMP with time for each dyes. All the values or data are represented in mean of three replications (n = 3) and bars indicated the standard error of mean.

### Sensitivity

The sensitivity of the LAMP was performed under optimized conditions using different template DNA concentrations and the results were visualised for each dyes. There were no positive results observed for all the dyes when the template was less than 10^−7^ ng from Xap1 isolate (Fig 5a). All the positive results from visualized LAMP assay was further assessed on 2% agarose gel and the band pattern was slightly faint with respect to decrease in template DNA concentration (Fig 5b). The similar dilutions were also tested for its sensitivity using PCR based amplification of Xap specific regions where the amplification was up to a template concentration of 10^−2^ ng (Fig 5c). We observed that the EtBr had highest fluorescent among the dyes used for the assay (Fig 5d).

**Figure 5.**
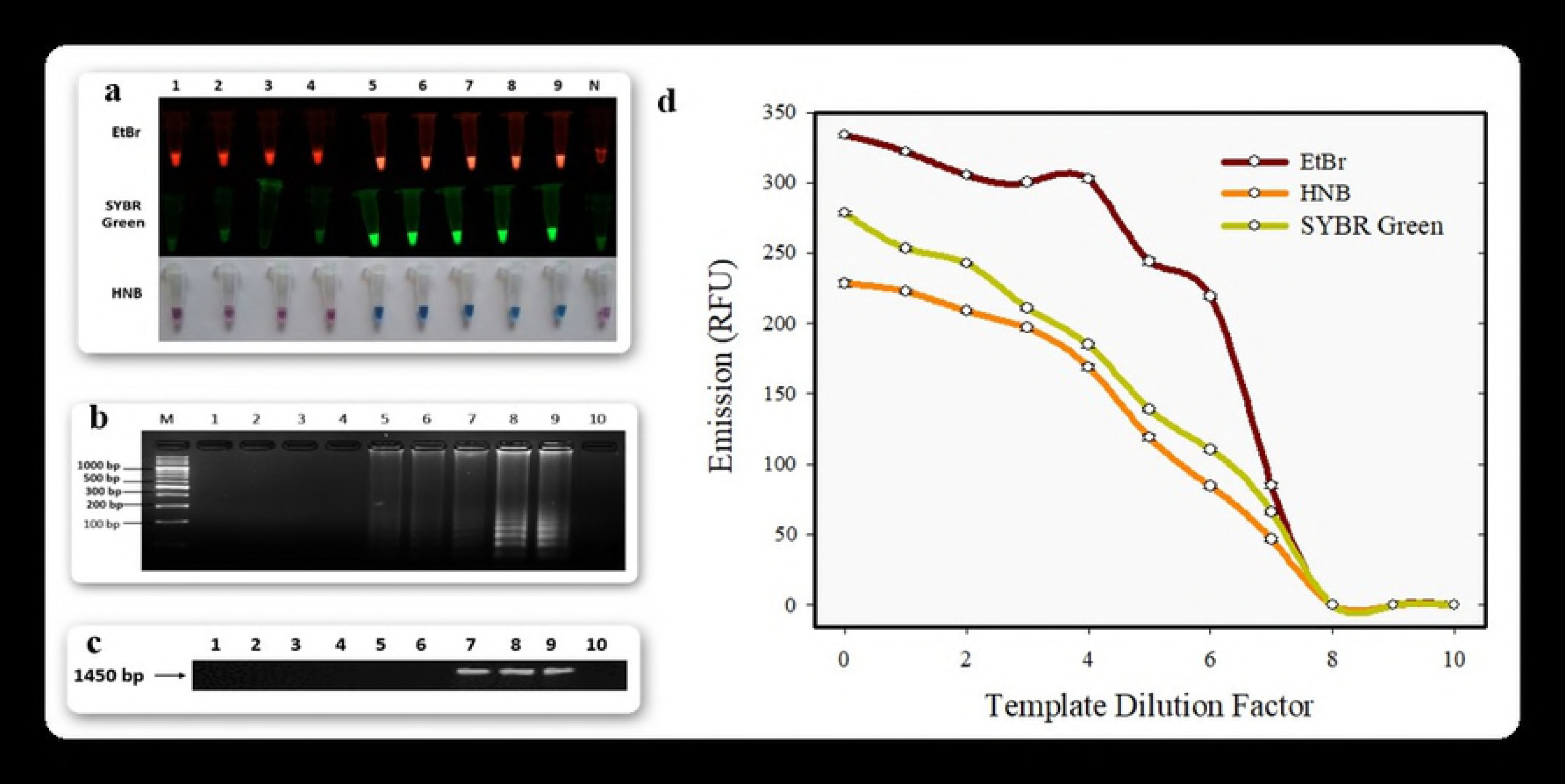
Sensitivity assay of LAMP and PCR methods. (**a**) Sensitivity of LAMP with dyes EtBr, SYBR Green and HNB ranging from 10 fold dilution of template 1.52 ng/μL for 10 times. (**b**) Sensitivity of LAMP was visualized in agarose gel, Lanes (1 to 9) indicated 10 fold serial dilutions and Lane N indicated negative control without any template. (**c**) Sensitivity of PCR was visualised in agarose gel with a product size of 1450 bp, Lanes (1 to 9) indicated 10 fold serial dilutions and Lane N indicated negative control without any template. (**d**) The sensitivity graph of the dyes were evaluated with 10 fold serial dilution from 1.53 ng/μL. All the values or data are represented in mean of three replications (n = 3) and bars indicated the standard error of mean.

### Specificity

Specificity of the LAMP assay was evaluated using 10 Xap and 29 non-Xap isolates (Table 4). These non-Xap bacterial isolates belonged to other bacterial species collected from different regions in Karnataka, India as detailed in Table 1. The DNA template obtained from Xap isolates which showed positive results when analyzed through the DNA intercalating dyes (Fig 6a). Further, it was reconfirmed on 2% agarose gel electrophoresis (Fig 6b) and the absorbance intensity for the dyes was validated (Fig 6d). The 16s rRNA was amplified to confirmed further for these Xap spp. (Fig 6C) for its specificity. To avoid any false positive results, 29 bacterial isolates collected from pomegranate were analysed which ultimately showed negative result for LAMP assay (Fig S1).

**Table 4.**
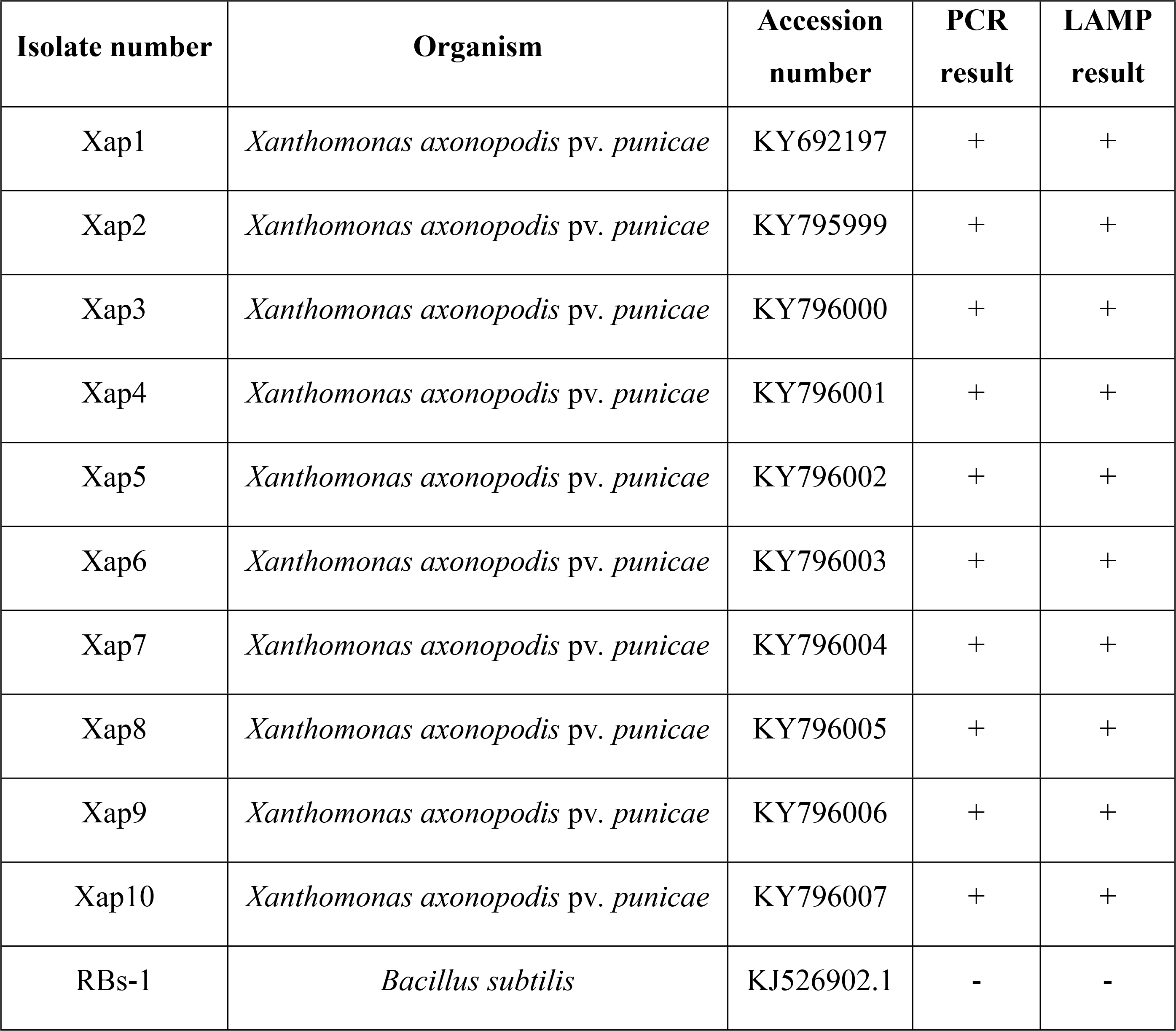
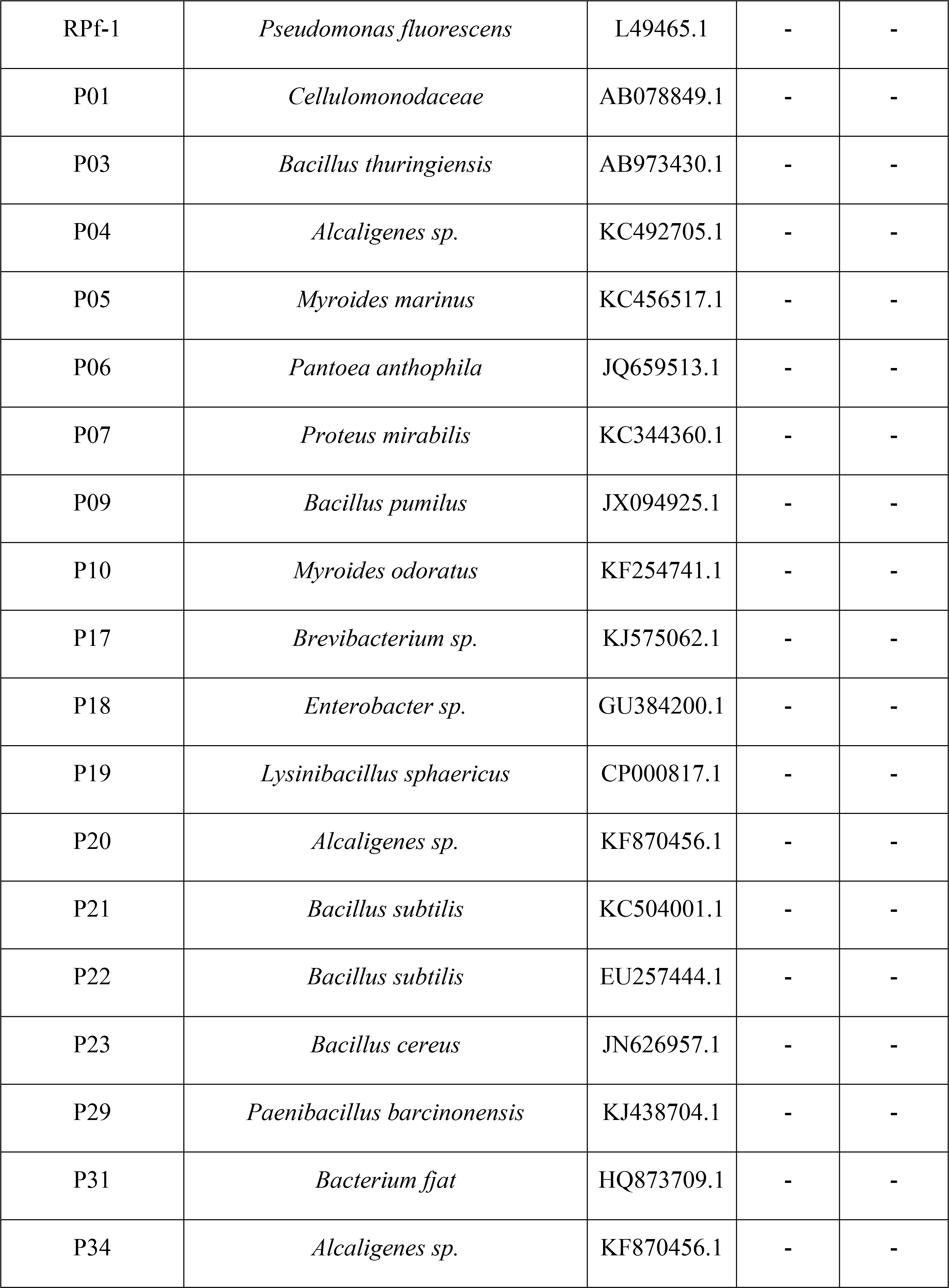
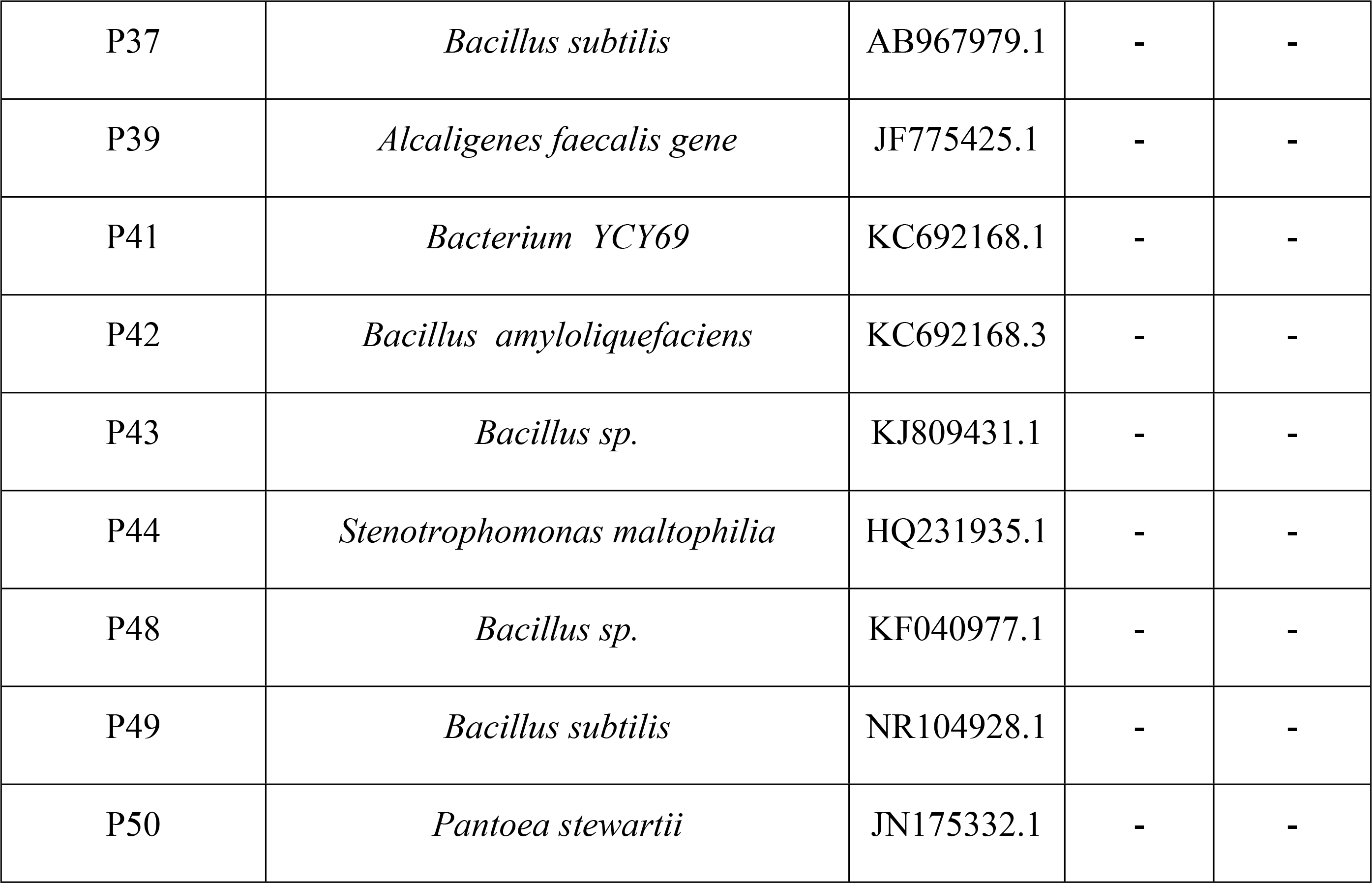
Details of Xap and non-Xap bacterial isolates used for specificity assay. All the collected isolates checked for amplification of Xap gene in both PCR and LAMP assays. The results of the PCR and LAMP assay are indicated as positive (+) or negative (−) based on amplification status using designed specific primers in PCR and also visualizing through three intercalating dyes with amplification for LAMP assays.

**Figure 6.**
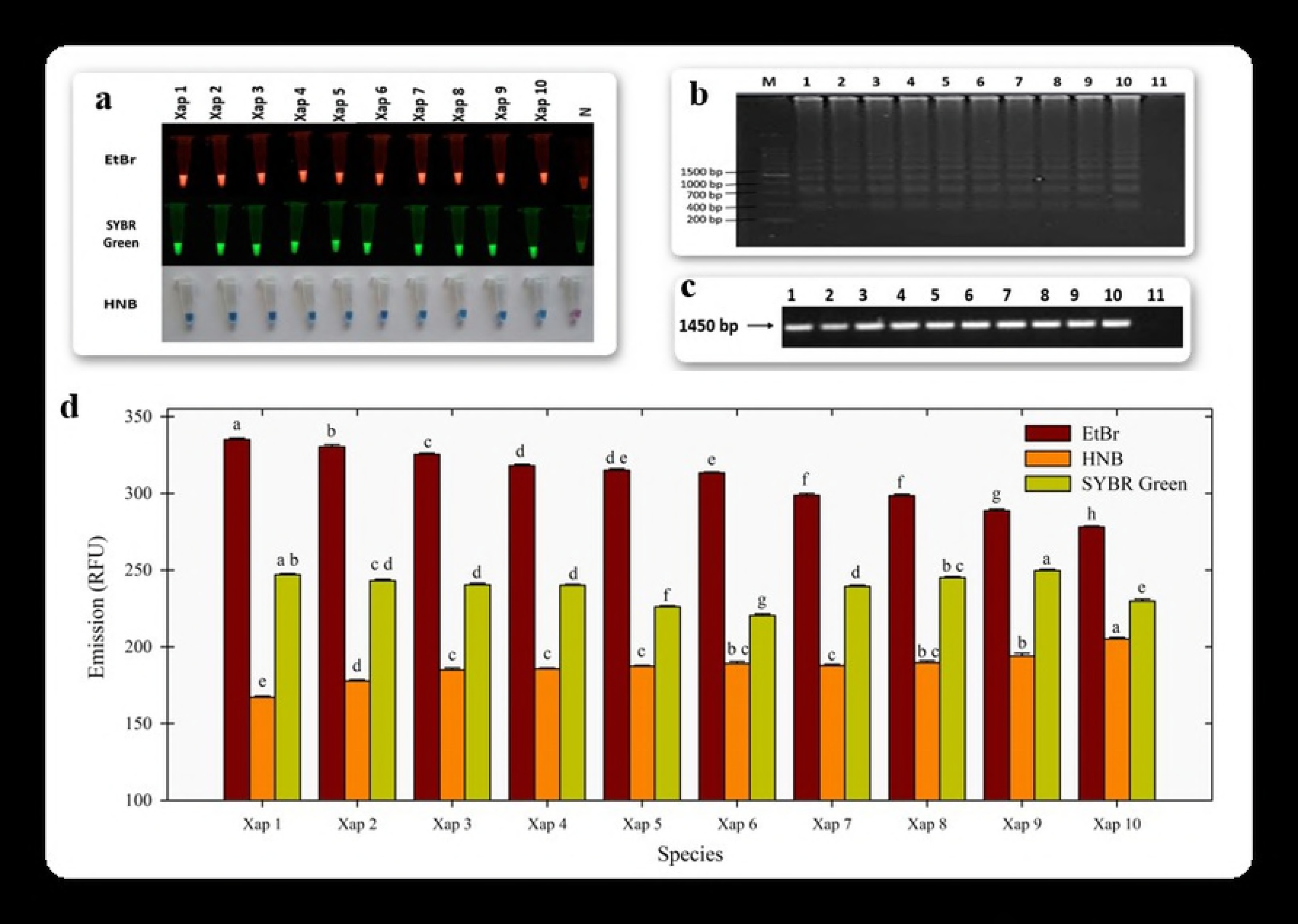
Specificity assay of LAMP and PCR methods. Ten isolates of *Xanthomonas axonopodis* pv. *punicae* under LAMP assay. (**a**) Accuracy of LAMP assay for the Xap isolates were validated using all the dyes, Xap 1 to Xap 10 in the Lane 1 to 10 and negative control in Lane N. (**b**) Specificity of LAMP was confirmed in 2% agarose gel, Lane 1 to 10 indicated Xap 1 - 10 isolates and Lane N indicated negative control without any template. (**c**) PCR products for the ten isolates was visualised by agarose gel, Lane 1 to 10 indicated Xap 1 - 10 isolates and Lane N indicated negative control without any template. (**d**) The specificity graph of LAMP for the 10 isolates of Xap tested against each dyes. The lowercase of the same letters which are indicated are not significant different between the isolates in any particular dye (Duncan’s multiple range test, P<0.05). All the values or data are represented in mean of three replications (n = 3) and bars indicated the standard error of mean.

### *In planta* detection

This study aimed to detect Xap by LAMP assay at the early crop stage where the leaf was mechanically pricked with bacterial blight pathogen to inoculate it to healthy pomegranate plants. Three months old pomegranate plant (Bhagwa va.) was grown under controlled environment in a glass house with 85% humidity. Xap1 bacterial culture was resuspended in sterile water and inoculated on healthy plants using pin prick method. The prominent symptoms were observed from the 11^th^ day after post inoculation, the oily lesions on the leaf surface were observed, which are characteristic symptoms of the bacterial blight disease. The crude DNA isolation method was followed to isolate DNA from the inoculated plants, every alternative day. After the pathogen infectivity in the healthy plants, LAMP assay in agarose gel showed positive from 7^th^ day onwards (Fig 7b) and for visualizing all the three dyes showed positive LAMP results from the 7^th^ day onwards (Fig 7a), in which the EtBr viewed to be most prominent comparing with other and its absorbance intensity was increased with days (Fig 7d). All the crude isolated DNA samples in PCR method showed amplification from 11^th^ day onwards (Fig 7c).

**Figure 7.**
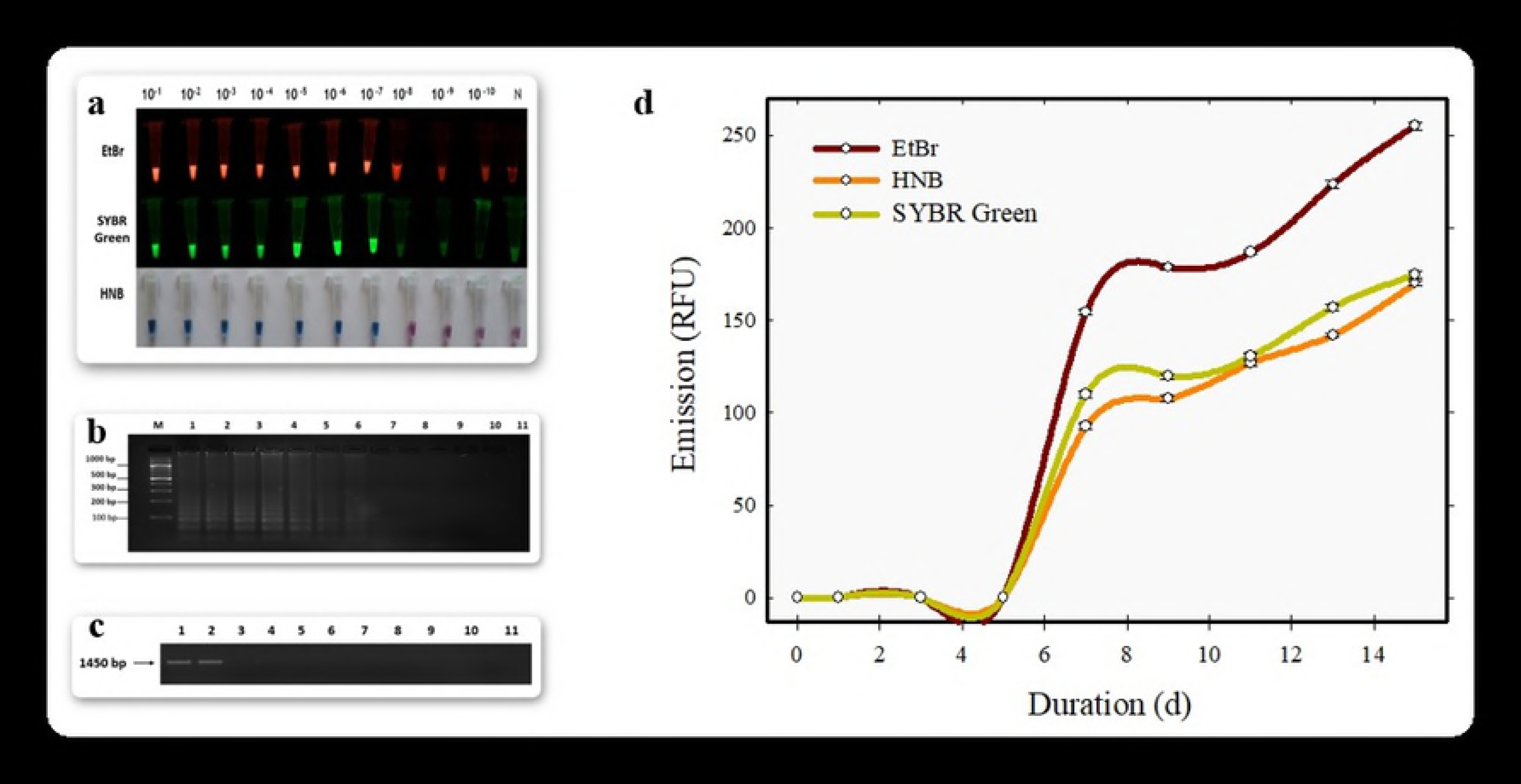
*In planta* validation studies carried out in glasshouse. (**a**) The crude samples were validated in LAMP and results were visualized through EtBr, SYBR Green and HNB dyes. (**b**) Glass house samples of LAMP was reconfirmed in agarose gel, Lanes 1 - 9 indicated (0, 1, 3, 5, 7, 9, 11, 13, 15 days respectively) and Lane 10 indicated negative control. (**c**) Glasshouse validation of PCR was visualised in agarose gel, Lanes (1 to 9) indicated (0, 1, 3, 5, 7, 9, 11, 13, 15 days respectively) and Lane N indicates negative control. (**d**) *In planta* validation studies plotted in graphs for the dyes. All the values or data are represented in mean of three replications (n = 3) and bars indicated the standard error of mean.

### Statistical data validation

The method used in LAMP assay with the dyes were validated with additional 10 Xap isolates. Duncan’s Multiple Range Test (DMRT) was used among the ten isolates for each dyes. Duncan’s Multiple Range test (DMRT) is a post hoc test to measure specific differences between pairs of means. The least error mean square was observed among the 10 isolates in SYBR Green followed by EtBr and HNB. The ‘t’ test was used within the isolates where all the dyes were significant at degree of freedom 29, EtBr (94.06), HNB (103.74) and SYBR Green (138.58) at 95% confidence level.

## DISCUSSIONS

This study aimed to show case the importance of LAMP in an early detection of pathogen. Early detection of plant pathogens could possibly result successful disease management for the crop. The spread of the pathogen namely *Xanthomonas axonopodis* pv. *punicae* is emerging as massive threat to the pomegranate cultivators, worldwide. The early detection of the pathogen (Xap) in pomegranate was attempted with LAMP assay which address a better alternative than the conventional PCR method, which are eventually highly reliable, cost-effective, very sensitive, more specific and rapid. The early detection would mainly benefit to diagnose and minimize the crop damage [25]. PCR based detection is highly reliable on high quality DNA with high quantity of template, typically 1 ng to 1000 ng [27]. The flow of our LAMP assay is diagrammatically described in Fig 8.

**Figure 8.**
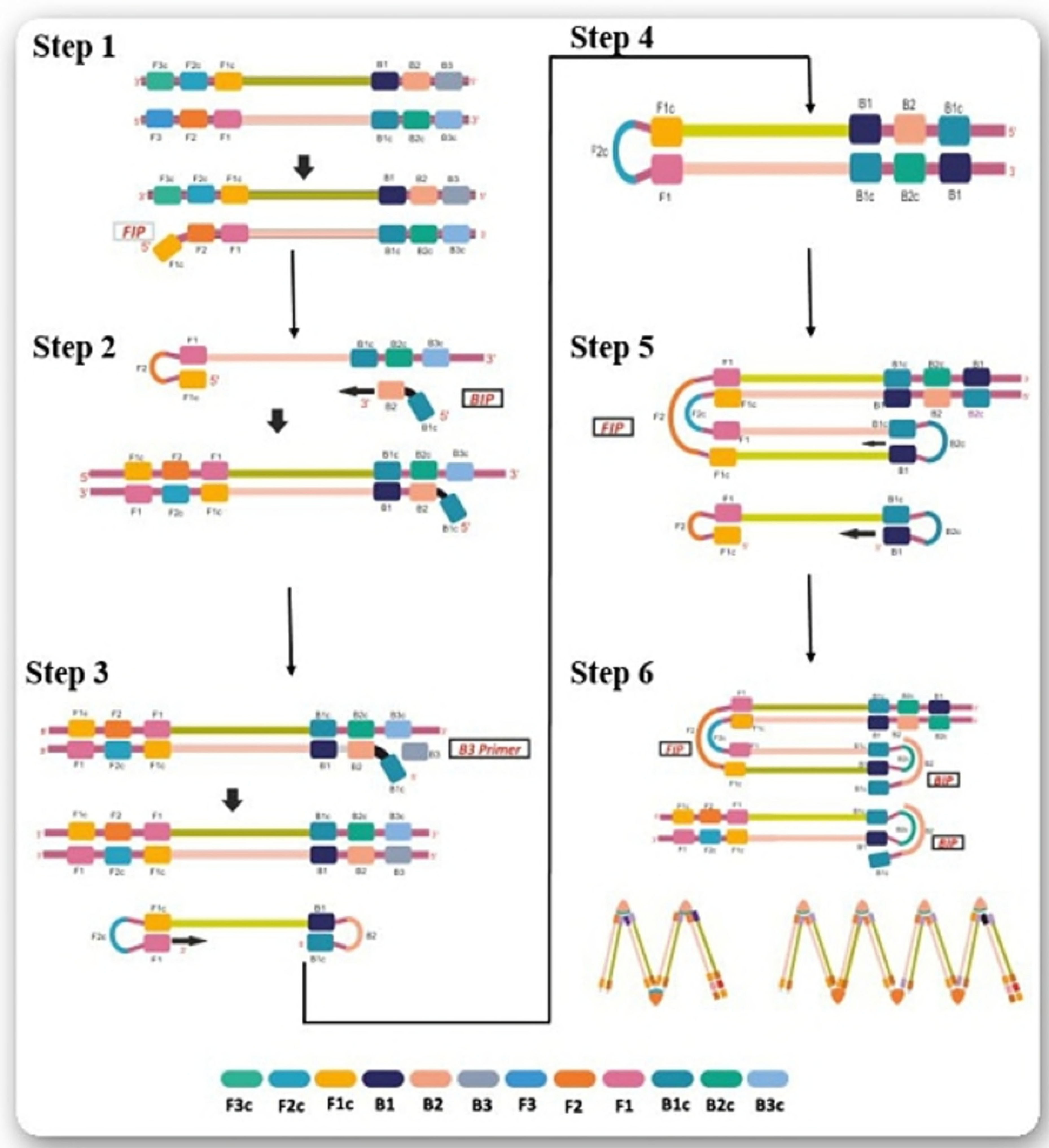
Overall workflow involved in the LAMP assay. Step **1**. The FIP region segregates into F2 region at the 3’end and F1c region at the 5’end.the F2 region hybridizes to F2c of the target and further initiates the synthesis of complementary strand. Step **2**. The loop formed at 5’ end acts as a template for BIP region. Once B2 hybridizes to B2c of the template, DNA synthesis is initiated resulting in the formation of complementary strand and opening of the 5’ end loop. Step **3**. Outer primer B3 hybridizes to B3c region of the target and elongates, resulting in the formation of a dumbbell shaped DNA. Step **4**. The dumbbell shaped DNA, now converts to a stem loop structure. The FIP hybridizes to form a loop of the stem-loop DNA structure to initiate LAMP cycling. Strand synthesis is started. The F1 strand is displaced to form a new loop at the 3’ end. Step **5**. The elongation takes place displacing the FIP strand, which once again forms a dumbbell, shaped DNA. Step **6**. The obtained products then serves as template for the BIP primed strand displacement reaction in the subsequent cycles. The final products obtained are a mixture of stem loop DNA with various stem lengths structures with multiple loops.

We have amplified Xap using Xap-specific primers with a minimum template concentration of 1.53 × 10^−2^ ng in PCR. While our LAMP assay amplification was recorded in agarose gel till a least concentration of 1.53 × 10^−7^ ng. Report on bacterial pathogen by Jun-hai *et al.*, [28] detected bacterial blight caused by *Xanthomonas axonopodis* pv. *dieffenbachiae* infecting Anthurium in LAMP which had sensitivity of 1 fg, which is 100 times less sensitive when compared to our detection by LAMP assay on Xap. Sensitivity was comparably high in our method as we have used LAMP primers designed from 16S rRNA sequence, which could detect the pathogen even with low amount of 10 copies of genetic material[29,30]. In our study, we have used 10 Xap isolates and all assays were positive with LAMP dyes. There could always be a minute chance that the PCR method could amplify from some non-specifically primed reactions [31], so with these criterias, we used 29 non-Xap isolates in LAMP assay which resulted negative and thus it turned out to be highly specific and reliable method to detect Xap samples.

DNA intercalating dyes resulted in detection of pathogens in LAMP assay without performing gel electrophoresis. Most reports in LAMP assay involve EtBr dye for visualization [32,33]. Even though, it is a highly hazardous and showed closely similar sensitivity when compared to most of the florescent dyes [34]. We used SYBR Green [35] which is less mutagenic when compared to EtBr[36]. Both resulted same sensitivity level but EtBr resulted twice more in florescence intensity. We also used HNB dye to visualize through naked eye, which showed the same sensitivity level for EtBr and SYBR Green but the fluorescence intensity was comparably low.

LAMP reaction produces massive amount of amplification as many as 10^9^-10^10^ copies of the target regions within an hour,[37] as a result cross contamination could be generated only when indicator dye was added after the reaction like EtBr and SYBR Green. The DNA intercalators are small molecules that can reversibly bind in between the adjacent base pairs of a double-stranded DNA. The intercalating dyes binds to the dsDNA causing the nucleic acid to become fluorescent and therefore readily detected under UV light. EtBr and SYBR Green are intercalating agents which binds between the nucleotide pairs and fluoresce when exposed to UV light.

LAMP is highly sensitive and negligible amount of target DNA can also be effectively detected from the colour change. Hydroxy napthol blue (HNB), which is a colorimetric indicator of calcium and alkaline earth metal ions. Since HNB is added in the LAMP reaction prior to the process, the colour gradually changes from violet to blue as the dNTP’s decreases as amplification takes place. So, using dyes like HNB prior to the preceding LAMP reaction tubes can be ideal to prevent a chance of wrong detection [38].

For field level application, remote laboratories may not be well equipped where LAMP detection technique can be employed in diagnosis of pathogens. This assay can even detect Xap in much earlier crop stage than the traditional PCR method. In our greenhouse study after Xap inoculation the LAMP assay detected Xap on 7^th^ day while, PCR method detected from 10^th^ day onwards. However, the prominent symptoms of Xap infection on the leaf in the glass house study appeared only after 10^th^ day of inoculation (Fig 7)

Improvised field applications such as working with crude extracts from plants with minimal requirement will enhance the importance of the assay for rapid and accurate detection of the pathogens. Unlike most of the methods which involve complete DNA purification for LAMP assay [39], we used crude extract of the leaf sample for detection of Xap. In conclusion, with crude extraction and safer dyes with minimum equipment, our study has shown a scope to develop LAMP isothermal kit which could be handy in the fields for early detection of the disease and to adopt appropriate management strategies.

## Author Contributions

Conceived and designed the experiments: MKP, CM. Performed the experiments: PBP. Analysed the data: KSN, PBP, GVB. Contributed reagents/materials/analysis tools: MEP, KP, CM. Wrote the manuscript: KSN, GVB, PBP. Edited the manuscript: HBM, KTR. All author read and approved the manuscript for publication. The authors declare that they have no conflict of interest in the publication.

## Acknowledgement

The research work was funded by RKVY (Rashtriya Krishi Vikas Yojana) project, University of Agricultural Sciences Bangalore (UASB). Our team would like to thank A. Satish (Dept. of Soil Science, UASB) for assisting in GIS mapping construction. We are grateful to Basavaraj (Dept. of Plant Pathology, UASB), Sankrappa (Dept. of Horticulture, Bagalkot), Vinay Kumar (Dept. of Soil Science, UASB) and Edwin Raj (Plant physiology and Biotechnology, UPASI Tea Research Institute) for their technical assistance.

## Additional Information

Competing financial interests: The authors declare no competing financial interests.

